# pAgo-induced DNA interference protects bacteria from invader DNA

**DOI:** 10.1101/2020.03.01.971358

**Authors:** Anton Kuzmenko, Anastasiya Oguienko, Daria Esyunina, Denis Yudin, Mayya Petrova, Alina Kudinova, Olga Maslova, Maria Ninova, Sergei Ryazansky, David Leach, Alexei A. Aravin, Andrey Kulbachinskiy

## Abstract

Members of the conserved Argonaute protein family use small RNA guides to find their mRNA targets to regulate gene expression and suppress mobile genetic elements in eukaryotes. Argonautes are also present in many bacterial and archaeal species ^1–3^. Unlike eukaryotic proteins, several studied prokaryotic Argonautes use small DNA guides to cleave DNA, a process dubbed DNA interference ^4–8^. However, the natural functions and targets of DNA interference are poorly understood and the mechanisms of DNA guide generation and target discrimination remain unknown. Here, we studied the *in vivo* activities of a bacterial Argonaute nuclease CbAgo and demonstrated that it induces cleavage of multicopy genetic elements, including plasmids, transposons and repetitive chromosomal loci. Generation of small DNA guides employed by CbAgo requires cooperation between its intrinsic endonuclease activity and the cellular double-strand break repair machinery. The mechanism of guide generation ensures that small DNA guides are enriched in sequences that target foreign DNA and endows CbAgo with capacity to eliminate plasmids and fight phage infection. Similar principles may underlie the specificity of self-nonself discrimination by diverse defense systems in prokaryotes.

---

Argonaute (Ago) proteins are a ubiquitous family of guide-dependent nucleases found in all three domains of life ^1,3,9,10^. Eukaryotic Ago proteins (eAgos) are RNA-dependent RNA nucleases that suppress gene expression by binding to their RNA targets in the cytoplasm or in the nucleus ^11,12^. Prokaryotic Agos (pAgos) are extremely diverse in comparison with eAgos and are randomly distributed among bacterial and archaeal lineages suggesting their active spread by horizontal gene transfer ^1–3^. In contrast to their eukaryotic counterparts, several studied pAgos were shown to act *in vitro* as nucleases with distinct specificity toward DNA targets ^4–6,8,13–17^. The majority of studied pAgos also prefer to use short single-stranded DNA molecules as guides for their endonuclease activity, however, the molecular mechanism of DNA guide generation in bacterial cells has remained poorly understood. Two pAgo proteins studied *in vivo* were shown to decrease plasmid content and transformation efficiency in bacterial cells ^7,18^, leading to the hypothesis that pAgos use DNA interference to protect the cells against foreign DNA. However, the ability of pAgos to fight genuine invaders such as phages was not demonstrated and the factors that could instruct pAgos for the recognition of foreign genetic elements have remained unknown.

Here, we investigate the mechanism of DNA interference by a recently characterized pAgo nuclease, CbAgo from a mesophilic bacterium *Clostridium butyricum*, which acts as an efficient DNA-guided DNA nuclease under ambient conditions *in vitro* ^4,5^. We demonstrate that in bacterial cells CbAgo preferentially targets multicopy genetic elements and induces DNA interference between homologous sequences, acting in cooperation with the cellular DNA repair machinery and allowing the host cells to eliminate invader DNA.

## CbAgo binds small DNAs from *Ter* sites

To study the cellular activities of CbAgo, we purified it from bacterial cells and analysed associated nucleic acids (Extended Data Fig. 1a). CbAgo was bound to small guide DNAs (smDNAs) of 14-23 nt, as confirmed by their sensitivity to DNase treatment and resistance to RNase (Extended Data Fig. 1a, b). Sequencing of CbAgo-bound smDNAs showed a large diversity of sequences with moderate AT-bias near the 5’-end and downstream of the site of target cleavage (Extended Data Fig. 1c), suggesting that their biogenesis might depend on melting of these DNA regions in the context of double-stranded DNA (Extended data Fig. 1e).

Chromosomal mapping of CbAgo-bound smDNAs revealed that they are distributed through the whole genome, with two large peaks at the region of replication termination (Fig. 1a, b). The smDNA hotspots are restricted by the *Ter* sites from the outside and are asymmetric, with the higher peak corresponding to *TerC* and the lower to *TerA*. This correlates with the shorter and longer lengths of the right and left replichores, respectively, and different frequencies of replication termination at *TerC* and *TerA* ^19^.

**Fig. 1.**
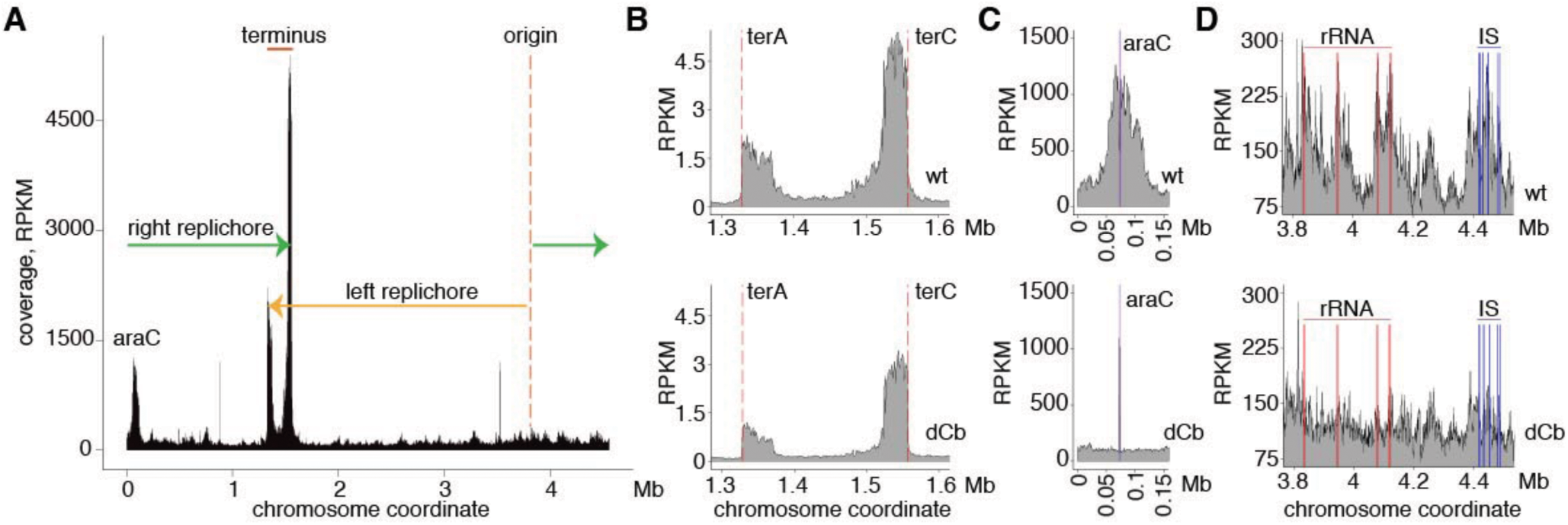
CbAgo targets specific genomic regions. (a) Distribution of smDNA guides associated with CbAgo along the bacterial chromosome. Replication origin and termination sites and the directions of replichores are indicated. (b) SmDNA peaks in the *Ter* region of the chromosome for wild-type (wt, top) and catalytically-impaired CbAgo (dCb, bottom). (c) SmDNA peak in the *ara* locus. (d) SmDNA profiles in the chromosomal region containing four rRNA operons (red) and a cluster of IS elements (blue). RPKM, density of smDNAs in reads per kilobase per million reads in the smDNA library.

To explore the relationship between replication termination and generation of smDNAs, we analysed CbAgo-bound nucleic acids in a strain lacking Tus, the protein that binds *Ter* sites and limits replisome progression. No smDNA peaks at *Ter* sites were observed in the *tus*^-^ strain. Instead, a single peak was located exactly opposite the origin of replication, close to *TerC*, likely corresponding to the main site of replisome collisions in this strain (Extended Data Fig. 2, *tus*^-^ strain). Therefore, replication termination by Tus-DNA complexes results in preferential smDNA generation at *Ter* sites.

## DNA interference between multicopy sites

When CbAgo was expressed from a plasmid with a region of homology to the chromosome, the *araC* gene, an additional strong peak of smDNAs appeared around the genomic *ara* locus (Fig. 1a, c). Small DNAs were generated as far as 30-40 kb from the *ara* locus indicating that the presence of the plasmid triggers smDNA processing at the identical genomic sequence that further spreads into flanking regions. This result suggests that CbAgo loaded with plasmid-derived guide DNAs attacks a homologous chromosomal region and induces DNA interference between the plasmid and chromosomal loci.

To confirm that CbAgo can be directed toward any chromosomal region when instructed with plasmid-derived guides, we analysed CbAgo-bound smDNAs in a strain encoding chromosomal CbAgo and containing a plasmid with the *lacI* gene. In this case, we observed a large peak of smDNAs around the chromosomal *lacI* locus and an additional peak around the second *lacI* gene present in this strain at the *attB* site (Extended data Fig. 2d). No *lacI* peak was observed in the absence of plasmid (Extended data Fig. 2c). CbAgo can therefore induce DNA interference between plasmid and chromosomal regions independently of its genetic location.

The observed processing of distant chromosomal loci by CbAgo loaded with plasmid-derived guides suggests a possibility of interference between other repetitive elements in the genome. Indeed, inspection of multicopy genomic regions revealed peaks of smDNAs around multimapping sequences, primarily ribosomal RNA (rRNA) operons and IS elements (Fig. 1d). The peak width in each case was in the range of dozen kilobases, far beyond the borders of ribosomal DNA (rDNA) and IS elements. These results demonstrate that CbAgo can target multicopy sequences of both plasmid and genomic origin and induce DNA interference between homologous regions.

## Role of the catalytic activity of CbAgo

To study the role of the nuclease activity of CbAgo in generation of smDNA guides, we analysed a catalytically impaired CbAgo variant, dCbAgo (catalytically <u>d</u>ead CbAgo) containing substitutions of key amino acids in the active site ^5^. When purified from bacterial cells, dCbAgo was loaded with smDNAs suggesting that its endonuclease activity is not strictly required to generate smDNA guides (Extended Data Fig. 1b). However, we observed significant changes in the smDNA profile along the genome in comparison with the wild-type protein. Preferential generation of smDNAs at multicopy sequences – *araC*, rDNA and IS elements – was completely abolished in the catalytically inactive dCbAgo mutant (Fig. 1c, d). Analysis of the ratio of smDNAs associated with WT and dCbAgo allowed precise mapping of the chromosomal regions whose targeting depended on the catalytic activity of CbAgo (Extended Data Fig. 3). This analysis confirmed the key role of CbAgo in smDNA biogenesis at the rDNA loci and IS elements, especially the multicopy families of IS1 and IS3. In particular, all copies of IS1 were found in the areas with the WT/dCbAgo ratio of >1, a dramatic enrichment in comparison with random sampling (p<0.0001).

In contrast to multicopy sequences, smDNA peaks were still observed for dCbAgo in the *Ter* region (Fig. 1b). However, smDNAs from this region were also enriched in the wild-type protein in comparison with dCbAgo (Extended Data Fig. 3). Therefore, the catalytic activity of CbAgo is indispensable for DNA interference between multicopy sequences and contributes to generation of smDNAs at the termination sites.

## Asymmetry in smDNA processing

Inspection of smDNA distribution in the genomic *ara* locus (undergoing DNA interference directed by the plasmid-encoded *araC* gene) or in rDNA loci (undergoing intrachromosomal DNA interference) revealed a striking asymmetry between the two DNA strands around the target gene. In both cases, the majority of smDNA reads corresponded to the DNA strands whose 3’-temini were oriented towards the target gene (Fig. 2b, WT CbAgo; Extended Data Fig. 4a). In the *Ter* region, most smDNAs were produced from the 3’-terminated DNA strand at each *Ter* site (Fig. 2c). The outer boundaries of smDNA peaks at *araC* and rDNA loci were defined by Chi sites oriented in the 5’→3’-direction in the preferentially targeted DNA strand (Fig. 2b). The inner boundaries of the *TerA* and *TerC* peaks of smDNAs corresponded to the first Chi site located in the 3’-terminated strand before the respective *Ter* site (Fig. 2c).

**Fig. 2.**
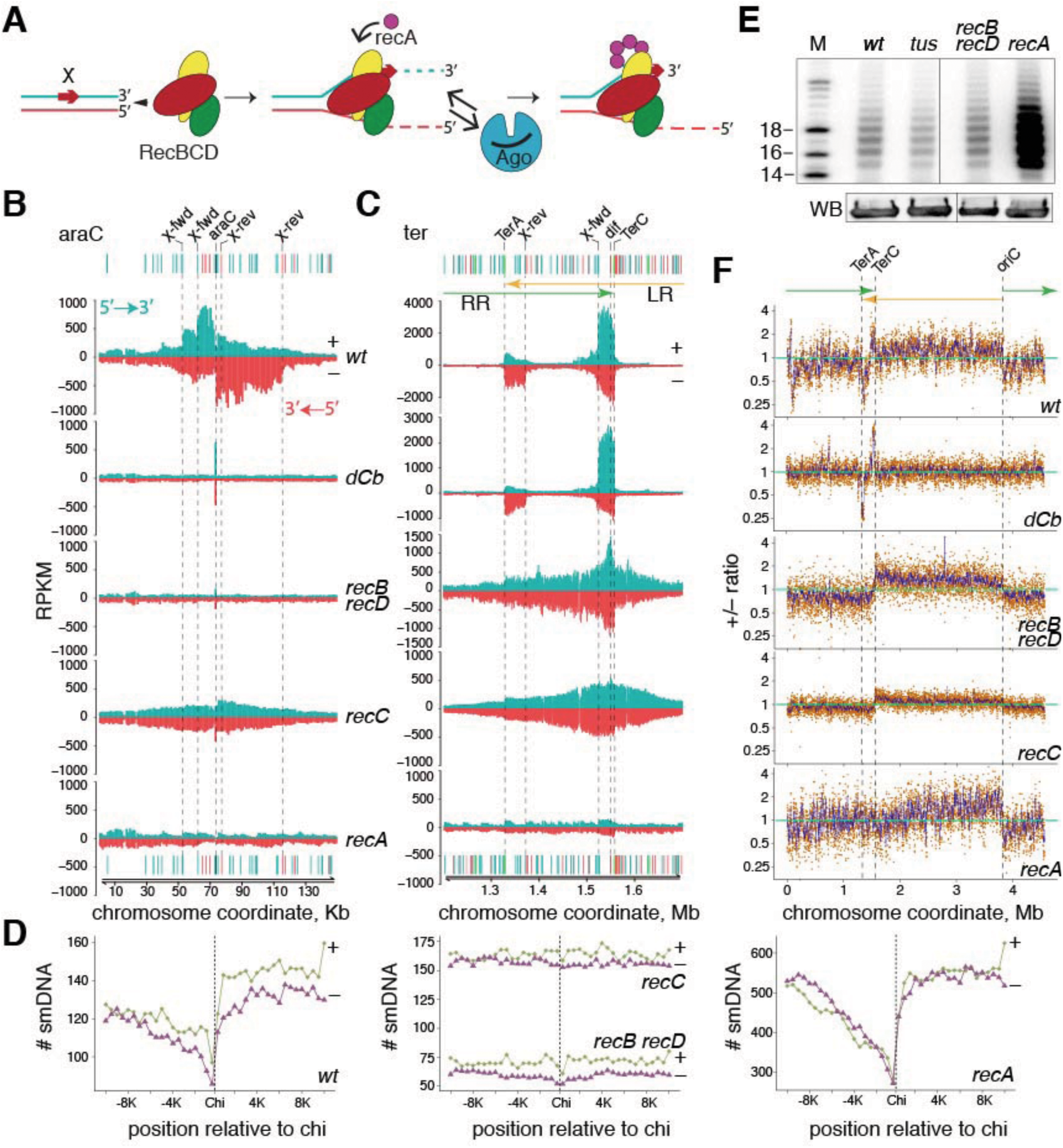
CbAgo cooperates with RecBCD in genomic DNA processing. (a) RecBCD determines the polarity of smDNA loading into CbAgo. (b) and (c) Strand-specific distribution of smDNAs in the *araC* locus (b) and *Ter* region (c) for strains with various genetic backgrounds. The reads from the plus and minus genomic strands are shown in green and red, respectively. Positions of Chi sites in surrounding genomic regions are indicated (forward for the plus strand and reverse for the minus strand); the closest Chi sites in the proper orientation are shown with dotted lines. LR and RR, leftward and rightward replichores. (d) Metaplot of the number of smDNAs around Chi sites in each genomic strand (red, plus-strand smDNAs for plus-strand Chi sites; green, minus-strand smDNAs for minus-strand Chi sites) in the 2-3 Mb genomic region. (e) CbAgo expression (WB, Western blotting) and smDNA loading into CbAgo in various bacterial strains. (f) Strand-specific asymmetry in smDNA distribution for various strains (ratio of RPKM values for the plus and minus genomic strands).

Chi sites are species-specific motifs (an 5’-GCTGGTGG-3’ octanucleotide in *E. coli*) recognized by the RecBCD or AddAB helicases-nucleases, which participate in double-strand break (DSB) repair and homologous recombination in various bacterial species ^20–22^. The RecBCD complex recognizes Chi in the 3’-terminated strand during DSB processing; this switches its activity from preferential cleavage of the 3’-terminated strand to preferential cleavage of the 5’-terminated strand thus generating a 3’-terminated single strand for RecA loading (Fig. 2a). The observed asymmetry of smDNAs and their dependence on Chi sites suggests that smDNA biogenesis is linked to formation of DSBs and their processing by RecBCD. More specifically, smDNA are preferentially generated from the 3’-terminated strand at the sites of DSBs. In the case of *araC* and multicopy elements, smDNAs are produced from both ends of the DSB presumably formed at the target loci. In the case of *Ter* sites, smDNAs are likely produced from double-stranded DNA ends, formed at *TerA* and *TerC*, by RecBCD moving in reverse direction relative to prior movement of the arrested replication forks.

To further explore the role of Chi sites in generation of smDNAs on the whole-genome level, we analysed smDNA distribution around Chi sequences across a 1 megabase region lacking strong smDNA peaks (from 2 to 3 Mb of genomic coordinates). This analysis demonstrated enrichment for smDNAs coming from the 3’-terminated strand at the 3’-side of Chi sequences, for Chi sites from both plus and minus genomic strands (Fig. 2d). Thus, generation of smDNAs is intimately linked to DSB processing and Chi site positioning in the genome.

## Role of the DSB repair machinery

To explore the role of the DSB repair machinery in smDNA biogenesis, we analysed strains with knockouts of individual components of the RecBCD complex (*recBrecD* and *recC*) and of RecA. The total amount of smDNA in CbAgo did not change significantly in the *recBrecD* and *recC* mutants (Fig. 2e). However, the polarity for smDNAs around Chi sites was eliminated in these strains, confirming that it is defined by RecBCD (Fig. 2d, *recBrecD* and *recC* strains).

No smDNA peaks at *araC* or rDNA loci were formed in the double *recBrecD* knockout strain demonstrating that RecB and/or RecD are essential for CbAgo-induced DNA interference (Fig. 2b and Extended data Fig. 2). The peaks were still present in the *recC* mutant but smDNA distribution became independent of Chi sites and symmetrical relative to the two DNA strands, because the RecBCD function was inactivated (Fig. 2b and Extended data Fig. 2). Therefore, different components of the RecBCD complex may have independent roles in DNA interference. In contrast, knockouts of not only *recB/recD* but also *recC* resulted in disappearance of the two smDNA peaks at *TerA* and *TerC*, with a single peak between them, suggesting that in wild-type cells the RecBCD complex orchestrates DNA processing at the *Ter* sites (Fig. 2c). In the mutant strains, the smDNA peak was independent of Chi sites and spread beyond the *Ter* sites. This likely indicates generation of DSBs in the terminus by septum closure during cell division with incomplete replication, previously observed for *recB* or *recC* mutants ^23,24^, followed by smDNA processing by CbAgo and/or other cellular nucleases.

The amount of smDNAs loaded into CbAgo was strongly increased in the *recA-*minus strain (Fig. 2e). This was accompanied by loss of specific enrichment at *ter*, *araC* or multicopy genomic loci. Instead, we noticed a shallow enrichment for smDNAs around the *ori* region, possibly reflecting the higher DNA content and/or a higher likelihood of DSB formation in this region (Fig. 2b and Extended Data Fig. 2). SmDNA distribution in the *recA* strain showed strong dependence on the Chi sites suggesting the involvement of RecBCD (Fig. 2d). Thus, in wild-type cells RecA guards the genome from excessive processing by RecBCD and CbAgo.

Analysis of the ratio of smDNAs originating from the two genomic strands on the whole-genome scale revealed a small but significant bias toward DNA strands whose 3’-termini were oriented opposite to the direction of each replichore, indicating that smDNAs may preferentially originate from the lagging DNA strand during replication (Fig. 2f). A similar bias was observed in the *recBrecD, recC* and *recA* mutant strains but not in wild-type cells expressing catalytically inactive dCbAgo (Fig. 2f). This additional asymmetry is therefore determined by the catalytic activity of CbAgo, suggesting that it can target the discontinuous lagging DNA strand during replication independently of RecBCD.

## DSB processing by CbAgo

The link between CbAgo-bound smDNAs and RecBCD suggests that DSBs might serve as a source for smDNA processing. To directly test the role of DSBs in smDNA biogenesis, we analysed CbAgo-associated smDNAs in bacterial strains with engineered DSBs, induced by either expression of the I-SceI meganuclease, recognizing its respective site in the genome^25^, or by a long palindrome (*pal*) processed by the host SbcCD (Mre11-Rad50) complex^26^ (both inserted in the *lac* locus) (Fig. 3a). In both cases, we observed highly efficient loading of smDNAs from the sites of breaks into CbAgo, with the peak size greatly exceeding the peaks at the *Ter* sites (Fig. 3b and Extended Data Fig. 4b). The size of the smDNA peak was smaller in a strain with a mutated I-SceI site, which is cleaved less efficiently^25^, demonstrating that smDNA production depends on the efficiency of DSB formation (Fig. 3b). The presence of the DSB also shifted the ratio between the *TerA* and *TerC* peaks in favor of *TerA*, likely as a result of impediment of the clockwise replisome, moving toward *TerC*, by the DSB formation (Fig. 3b). The regions flanking DSBs are therefore preferential substrates for generation of smDNA guides for loading into CbAgo.

**Fig. 3.**
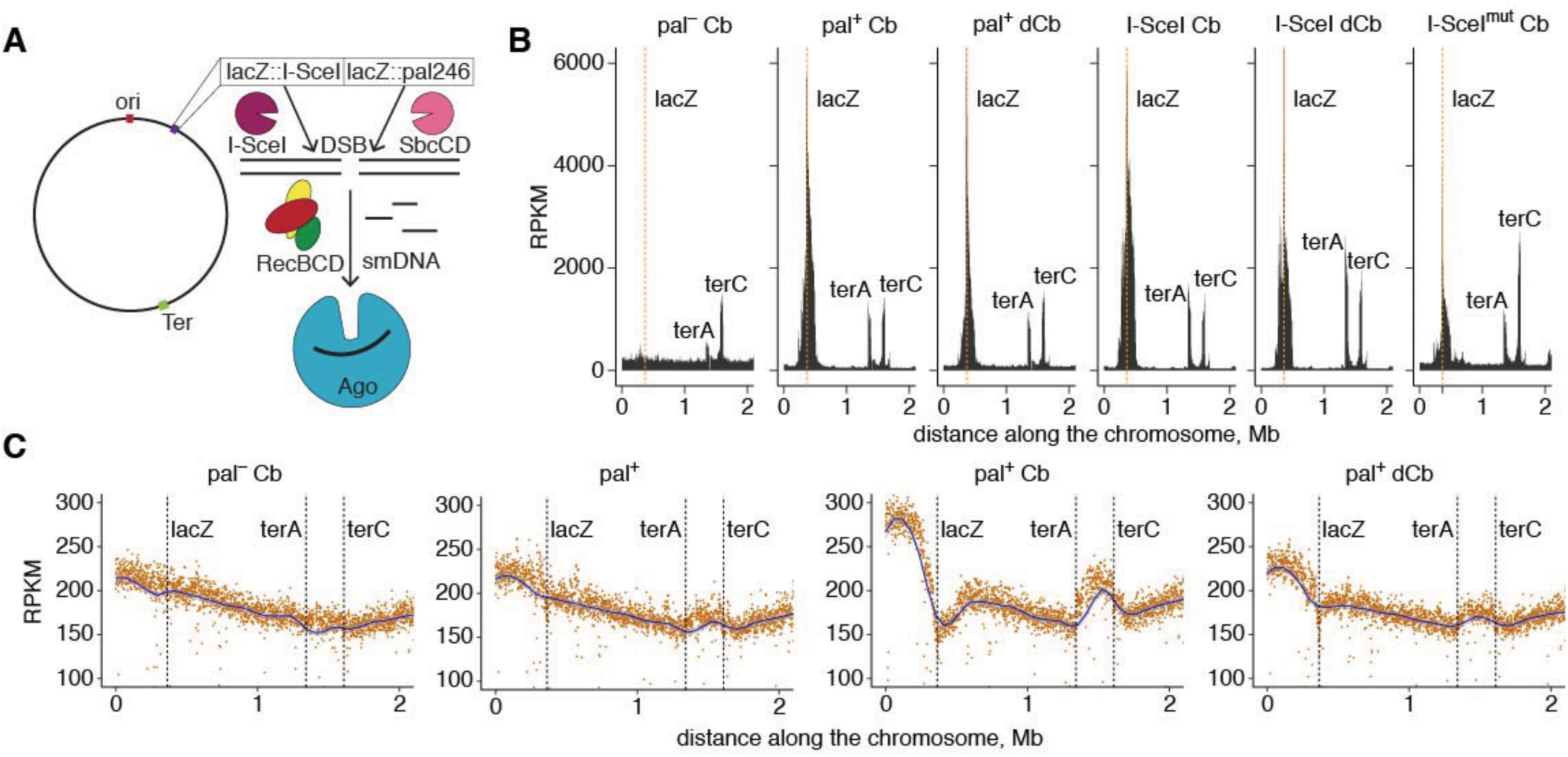
CbAgo attacks DSBs. (a) Formation of DSBs at the *lacZ* locus. (b) SmDNA abundance in the chromosomal region spanning the engineered DSBs (palindrome or I-SceI-dependent; I-SceI^mut^, the mutated cleavage site) and *Ter* sites. (c) Genomic DNA coverage in the same region in the control (*pal*^−^) and palindrome-containing strains depending on the expression of active CbAgo or dCbAgo. Positions of the *lacZ* locus and *Ter* sites are indicated.

Inactive dCbAgo was also loaded with smDNAs at DSBs, although with a somewhat lower efficiency than wild-type CbAgo (Fig. 3b). In the case of both WT and dCbAgo processing of engineered DSB involves RecBCD since the boundaries of the smDNA peaks are defined by Chi sites, with more smDNAs being produced from the 3’-terminated strand at each end of the DSB (Extended Data Fig. 4b). This suggests that although the catalytic activity of CbAgo is essential for DSB formation during DNA interference between multicopy elements, it is not absolutely required for RecBCD-dependent processing of preformed DSBs.

To test if generation of smDNAs at DSBs affects genomic DNA integrity, we used high-throughput sequencing to analyse DNA abundance along the genome. As expected, in wild-type cells the DNA abundance gradually decreases towards termination sites (Fig. 3c, *pal*-minus strain). No obvious decrease in the genomic DNA content is observed at the site of palindrome-induced DSB in the absence of CbAgo, with small overreplication at the site of termination, indicating that DSBs resulting from palindrome processing are rapidly repaired (Fig. 3c, *pal*^+^, and Ref. ^27^). Similarly, only a small decrease in the genomic DNA content at the palindrome site is observed in a strain expressing dCbAgo (Fig. 3c, *pal*^+^ dCb), despite it being loaded with smDNAs corresponding to this region (Fig. 3b). In contrast, expression of CbAgo strongly decreases DNA coverage at the site of engineered DSB, with increased overreplication in the *Ter* region (Fig. 3c, *pal*^+^ Cb). In the case of the permanent DSB introduced by the I-SceI expression, the genomic DNA content at the target site is decreased even in the absence of CbAgo, but expression of CbAgo further stimulates DNA degradation in this region (Extended Data Fig. 5). Therefore, CbAgo activity triggers massive DNA loss at DSBs.

## CbAgo fights plasmids and phages

Our results indicate that smDNA guides loaded in CbAgo are preferentially generated from multicopy sequences and the sites of DSBs. These observations suggest that CbAgo may attack mobile genetic elements, such as transposons, plasmids and phages, which have multicopy nature and often form free DNA ends in their life-cycle. Furthermore, these elements usually lack Chi sites that could prevent their processing by the cellular DSB repair machinery. Indeed, analysis of the distribution of CbAgo-bound smDNAs between different plasmids and the genome revealed that a disproportionally large fraction of smDNA reads (up to 20%) are derived from plasmid DNA and can be mapped to both plasmid strands (Fig. 4a, c). Even after normalization for the plasmid copy number, the fraction of plasmid-derived smDNA was one-order of magnitude higher than expected in the case of random sampling of cellular DNA, indicating that CbAgo preferentially uses plasmid sequences as guide DNAs.

**Fig. 4.**
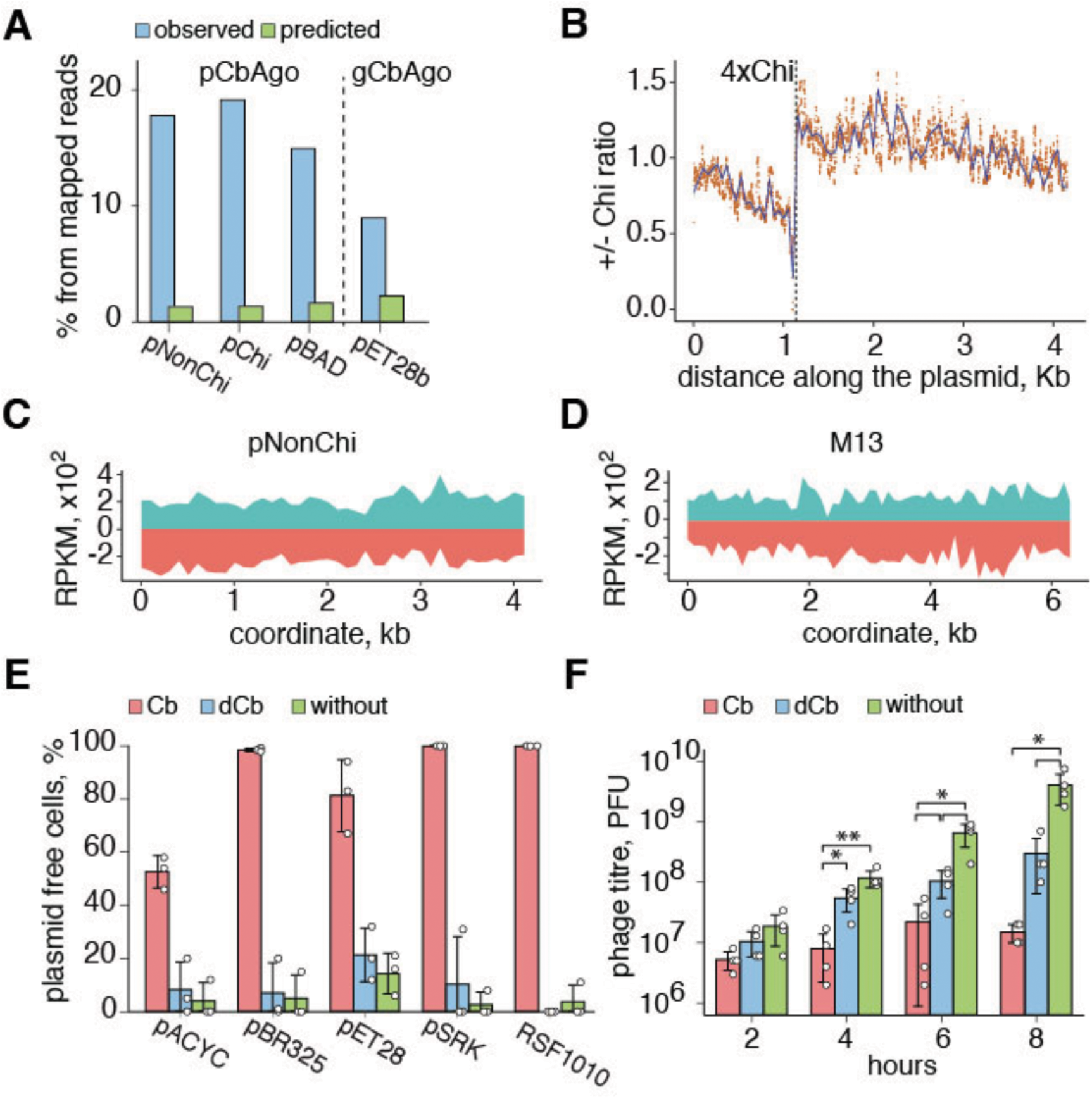
CbAgo fights plasmids and phage infection. (a) Enrichment of CbAgo with plasmid-derived smDNAs. The observed (blue) and expected (green) read numbers, based on random sampling of the genomic and plasmid DNA with normalization for the plasmid copy number (pCbAgo, plasmid-encoded CbAgo; gCbAgo, genomic CbAgo). pNonChi and pChi are pSRK derivatives without regions of homology to the genome; pChi contains four tandem Chi sites. (b) Asymmetry in smDNA distribution around the plasmid Chi sites (the ratio of smDNAs from the plus plasmid strand, containing Chi in the proper orientation, for the pChi and pNonChi plasmids). (c) Distribution of smDNAs along the plasmid sequence. (d) Distribution of smDNAs along the M13 genome. The reads from the plus and minus DNA strands are shown green and red, respectively. (e) CbAgo-induced loss of plasmids. The number of plasmid-free cells was measured after 5-9 passages in the absence of selective markers. Means and standard deviations from three independent measurements. (f) Decrease in M13 titers depending on the expression of CbAgo or dCbAgo. Means and standard deviations from four independent measurements (*p<0.05).

To reveal whether RecBCD participates in plasmid processing by CbAgo, we compared two plasmids with identical sequences, except for four tandem Chi sites present in one of them. The overall amount of plasmid-mapping smDNAs was similar between the two plasmids (Fig. 4a), probably because Chi sites at a single location could not prevent plasmid degradation. However, the smDNA distribution had a clear bias for smDNAs being produced from the 3’-side of the Chi sites, suggesting the involvement of RecBCD in their processing (Fig. 4b).

We then tested the effect of CbAgo expression on plasmid maintenance in bacterial cells. In the absence of selection for plasmid genes, control strains lacking CbAgo fully maintained the plasmids (Fig. 4e). In contrast, expression of wild-type, but not inactive CbAgo, led to fast and quantitative loss of plasmids from various incompatibility groups after few passages (Fig. 4e).

Next, we tested the effect of CbAgo on infection of bacterial cells with phage M13. Sequencing of CbAgo-associated smDNAs isolated from the infected strain revealed a considerable number of M13-derived smDNAs, which mapped to both strands of the circular phage genome, indicating that pAgo is loaded with smDNAs during phage infection (Fig. 4d). Expression of wild-type CbAgo reduced phage titers from 15 to 270-fold at 4 to 8 hours past infection (Fig. 4e). Expression of dCbAgo had a smaller but significant effect on phage titers (13-fold decrease at 8 hours). Therefore, CbAgo can protect the cells from foreign genetic elements, plasmids or phages.

## Discussion

Our results show that foreign DNA – plasmids, transposons and phages – is a preferential substrate for processing by CbAgo and, possibly, other DNA-targeting pAgo nucleases. CbAgo induces DNA interference between multicopy sequences present in the cell that likely underlies its functional activity against invading genetic elements. CbAgo cooperates with the cellular DNA break repair machinery and attacks vulnerable DNA regions containing breaks and free DNA ends that are formed during DNA replication, repair, and interference. In this model (Extended Data Fig. 6), preferential targeting of foreign DNA by CbAgo can be explained by: (i) the higher copy number of invader DNA thus allowing the loading of multiple guides into pAgo and resulting in more efficient DNA targeting over unique genomic regions; (ii) the ability of pAgo to target replication intermediates, which are located in smaller and therefore more vulnerable replicons in plasmids and phages; (iii) the relative absence of Chi sites in foreign DNA, thus making it a better substrate for RecBCD/AddAB processing^28^. Loading of locus-specific guides into pAgo induces DNA interference and processing of new smDNAs from the target locus, leading to amplification of guide DNAs and effective DNA destruction (Extended Data Fig. 6).

CbAgo targets specific genomic regions, including multicopy loci, the sites of pre-existing DSBs and the *Ter* region of the chromosome. Thus, in addition to fighting extra-genome invaders such as plasmids and phages, CbAgo-dependent DNA processing may potentially play roles in other genomic processes such as elimination of repetitive genetic elements, DNA repair and recombination, response to DNA damage and suicidal systems ^10^. CbAgo might also participate in the resolution of DNA intermediates during replication termination, as was recently proposed for another active pAgo nuclease, TtAgo from *Thermus thermophilus* ^29^.

pAgo proteins are randomly distributed among bacteria and archaea and their phylogeny does not correspond to the phylogeny of host species, as a result of horizontal gene transfer ^1–3^. pAgos may therefore get adapted to cooperate with various types of DNA processing and recombination machineries. Analysis of genomic associations of pAgos with cellular DSB repair machineries shows diverse combinations of factors in various bacterial species, with a substantial fraction of catalytically active pAgos found in the same genomes with AddAB or RecBCD (Extended Data Fig. 7). As we demonstrated, individual components of the RecBCD complex can also have independent functions in DNA interference. On the other hand, RecA protects the genome from indiscriminate degradation by CbAgo and RecBCD.

The finding that pAgos use DNA guides to protect bacteria against invaders highlights the conserved function of Argonaute-mediated nucleic acid interference (RNAi and DNAi) as a universal defense system that has survived billions years of evolution in both prokaryotes and eukaryotes ^30^. Prokaryotes have another defense system, CRISPR-Cas, whose activity is also based on complementary recognition of target genetic elements, but that uses different components and biochemistry. Remarkably, both pAgo and CRISPR-Cas coopt the cellular DNA break repair machinery to generate guide nucleic acids for their distinct effector complexes. Spacer acquisition by type I and II CRISPR-Cas systems was shown to be dependent on DNA replication and the RecBCD/AddAB machinery resulting in specific targeting of the *Ter* region and DSBs ^31,32^. Thus, pAgo and CRISPR-Cas may rely on similar principles for differentiation between ‘self’ and ‘nonself’ genetic elements. Similarly to Cas nucleases, pAgos can potentially be used for genomics applications, in particular as an instrument to study the genome architecture and DNA processing in connection with DNA replication, recombination, repair and transcription.

## Methods

### Plasmids and strains

Plasmids and bacterial strains used in this study are listed in Supplementary Tables S1 and S2. All strains are isogenic to either *E. coli* BL21(DE3), MG1655 or BW27784. *E. coli* cells were routinely cultivated in standard LB Miller broth (2% tryptone, 0.5% yeast extract, 1% NaCl, pH 7.0) with the addition of appropriate antibiotics (ampicillin, 100 μg/ml; kanamycin, 50 μg/ml; chloramphenicol, 12.5 μg/ml). All plasmids were introduced into recipient strains using a standard electroporation protocol according to the manufacturer’s instructions (BioRad Gene Pulser Xcell, 2.5 kV, 0.2 cm cuvettes).

For the construction of the pChi and pNonChi plasmids, a tandem of four consecutive Chi-sites was inserted into the pSRKKm plasmid via the Gibson assembly technique. Briefly, 5 μM of oligonucleotides 4xChi_fwd and 4xChi_rev (or 4xnonChi_fwd and 4xnonChi_rev) (Table S3) were mixed together in 50 μl of 1^x^ annealing buffer (10 mM Tris-HCl, 50 mM NaCl, 1 mM EDTA, pH 8.0), incubated at 98 °C for 5 minutes, slowly cooled down to 25 °C in a thermocycler, and cloned into pSRKKm linearized at the NheI and EcoRI sites, in order to remove a 1661 bp fragment containing regions of homology with the *E. coli* chromosome (*lacI*, *lacZ*⍺).

*E. coli* knock-out strains were obtained via Red-mediated gene disruption method essentially as described in ^33^. Briefly, kanamycin- or chloramphenicol-resistance cassettes were amplified with primers that contain 45 nt homology arms to the genomic regions flanking the target ORF from pKD4 or pKD3 plasmids, respectively, gel-purified and transformed into electrocompetent BL21(DE3) cells expressing the lambda-RED recombinase system under the control of the araBAD inducible promoter from the pKD46 plasmid. After 4 hours of recovery, the cells were plated onto the selective LB plates. The mutant alleles with selective markers were then transferred to parent strains by P1 transduction. The correct genomic integration events were verified by PCR and sequencing. The absence of the expression of the B, C and D subunits of the RecBCD complex was additionally checked by Western blotting with anti-RecB, anti-RecD and anti-RecC rabbit antisera (kindly provided by Dr. Gerry Smith).

*E. coli* strains with chromosomal insertions of the CbAgo gene were obtained starting from the MG1655Z1 strain containing a Z1 cassette with spectinomycin resistance (*LacI, TetR, SpR*) in the chromosomal *attB* site ^34^. MG1655Z1::CbAgo was constructed by insertion of a PCR product disrupting the Sp^R^ gene inside the Z1 cassette and containing two strong transcription terminators (t0 and BBaT1006), N-His6 CbAgo under the control of TetP promoter, BBaT1001 terminator and the *cat* gene. Positive Sm^S^ Cm^R^ clones were selected and verified by PCR and sequencing. This strain was used as a donor strain for P1 transduction of the wild-type MG1655Z1, NEB Turbo and strains containing the palindrome and I-SceI cleavage sites inside *lacZ*. The I-SceI cleavage site (TAGGGATAACAGGGTAAT) was present in the *lacZ* locus of the chromosome in the *E. coli* strain DL2917 ^25^. The mutant variant of the cleavage site I-SceI_mut_ in strain DL4977 contained two single-nucleotide substitutions relative to the wild-type sequence (T<u>T</u>GGGATAACAGGGTAA<u>A</u>, underlined). Strain DL2859 contained a 246 bp palindrome in *lacZ* and strain DL1777 was used as a control with the same genetic background ^26^. Strains carrying dCbAgo were constructed using the same protocol.

### Culture conditions for CbAgo expression and smDNA library preparation

In the case of plasmid-encoded CbAgo, the cells were transformed with plasmids encoding wild-type CbAgo or dCbAgo (pBAD/His B_CbAgo and pBAD/His B_dCbAgo, Table S1). After overnight growth on LB plates, a single colony was transferred into 5 ml of LB media supplemented with appropriate antibiotics and 0.2% glucose and grown overnight at 37°C. The cells were inoculated into 0.5 L of fresh LB supplemented with 0.2% glucose and appropriate antibiotics and aerated at 37°C until OD_600_ = 0.3-0.4. At this point, the temperature was adjusted to 30°C and after 20 min CbAgo expression was induced by the addition of L-arabinose to 0.01% for 3 hours. In the case of genomically-encoded CbAgo the cells were inoculated into LB medium that already contained the inducing agent (anhydrotetracycline, 200 ng/ml) and allowed to grow at 24°C until OD_600_ = 1.0. The cells were collected by centrifugation at 7000 g, 4°C for 15 minutes and kept frozen at −20°C.

### Isolation of smDNAs

All purification steps were carried out at 4°C. The cell pellet was resuspended in lysis buffer (50 mM Tris-HCl, 250 mM NaCl, 5% glycerol, 0.5 mM β-mercaptoethanol, pH 7.4 at 23°C) supplemented with EDTA-free protease inhibitor cocktail (Roche) at approximately 7 ml per gram of wet weight and subjected to 3 rounds of disruption on a high-pressure homogenizer (Avestin) at 12000 psi. The cell lysate was clarified for 30 min at 35,000 g. CbAgo (dCbAgo) was pulled down on Talon metal affinity resin (Takara) charged with Co^2+^ for 2 hours at 4°C (300 μl of bead suspension per sample) with constant rocking. The beads were collected by centrifugation (300 g, 3 min) and washed one time with lysis buffer and three times with lysis buffer supplemented with 10 mM imidazole (140 ml in total). CbAgo-smDNA complexes were eluted in 500 μl of lysis buffer supplemented with 300 mM imidazole.

Nucleic acids associated with CbAgo were extracted with phenol:chloroform:isoamyl alcohol (PCI) mixture (25:24:1, pH 8.0) according to a standard procedure. Briefly, the protein sample was mixed with an equal volume of PCI, vortexed intensively for 30 seconds and centrifuged at 13000 g for 3 min. The upper aqueous phase was carefully collected and treated twice with an equal volume of chloroform to eliminate any remaining traces of phenol. Nucleic acids were precipitated with three volumes of ethanol in the presence of a co-precipitating agent (PINK, Bioline) and dissolved in 20 μl of milliQ-grade water.

To determine the type of associated nucleic acids, the samples were treated with alkaline phosphatase (rSAP, NEB) at 37°C for 10 min with subsequent enzyme inactivation at 75°C for 7 min. The nucleic acids were subsequently radiolabeled with γ^32^P-ATP using T4 polynucleotide kinase (NEB) according to the manufacturer’s instructions. The samples were then divided into three equal parts and each part was treated with either DNase I (ThermoFisher), or RNase A (ThermoFisher) or left untreated. Radiolabeled nucleic acids were resolved on 19% denaturing polyacrylamide-urea gel and visualized on a Typhoon FLA 9500 imager (GE healthcare).

### Western blotting

The level of CbAgo expression and the amount of CbAgo used for smDNA purification was determined by Western blotting. Protein samples were mixed with 2x Laemmli sample buffer (120 mM Tris-HCl, 4% SDS, 4% β-mercaptoethanol, 10% Glycerol, pH 6.8) and heated at 95°C for 5 min, and then resolved by electrophoresis in a 4-20% Tris-glycine gel (BioRad). Proteins were transferred onto a nitrocellulose membrane in Towbin transfer buffer (25 mM Tris, 192 mM glycine, 20% methanol) using semi-dry procedure at 25 V, 1 A for 30 min (BioRad Trans-Blot Turbo). The transfer membrane was air-dried and then washed in PBS (10 mM phosphate buffer, 137 mM NaCl, 2.7 mM KCl) for 5 min. The membrane was blocked with blocking buffer (PBS, Tween-20 0.1% (v/v), non-fat milk 5% (m/v)) for 30 min at room temperature, and then incubated with anti-6xHis monoclonal antibodies (1:1000, Sigma) for 1 h at room temperature. The membrane was washed 4 times with PBST buffer (PBS, Tween-20 0.1% (v/v)), and after that incubated with HRP-conjugated anti-mouse secondary antibodies (1:10000, Sigma) for 1 h at room temperature and washed again as described above. Antigen-antibody complexes were detected with Immobilon ECL Ultra Western HRP substrate (Millipore) on a Chemidoc XRS+ imager (BioRad).

### SmDNA library construction and sequencing

Libraries for high throughput sequencing of smDNAs were prepared according to the previously described bridged (splinted) ligation protocol ^18^. For the purpose of visualization, a 5 μl aliquot of the smDNA sample (1/4 of total volume) was dephosphorylated and radiolabeled as described above. Unlabeled DNA was mixed with the radiolabeled aliquot and 2^x^ urea sample buffer (8 M urea, 20 mM EDTA, 0.005% Bromophenol Blue, 0.005% Xylene Cyanol and resolved under denaturing conditions in a 15% polyacrylamide-urea gel. Gel bands corresponding to 14-25 nt smDNAs were crashed and DNA was eluted in 0.4 M NaCl solution overnight at 21 °C with constant agitation. After ethanol precipitation, DNA was resuspended in 20 μl of milliQ-grade water.

Illumina-compatible adaptors were ligated to both ends of smDNA. For this purpose, 8 μl of 5^x^ Rapid Ligation buffer (ThermoFisher), 2 μl of 100 μM 5’-adaptor, 2 μl of 100 μM of 3’-linker, 2 μl of 100 μM oligonucleotide bridge 1, 2 μl of 100 μM oligonucleotide bridge 2 and 2 units of T4 PNK (ThermoFisher) were added to 20 μl of the purified smDNA solution. The reaction mixture was incubated overnight at room temperature. Ligated DNA fragments were recovered by 10% denaturing PAGE. The libraries were amplified and indexed according to the standard protocol (small RNA sequencing kit, NEB), except that PCR conditions were adjusted in order to prevent library over amplification. A series of test PCR were performed to optimize the amount of adaptor-ligated smDNA to achieve the desired amplification level in 3-4 cycles. smDNA libraries were sequenced using the HiSeq2500 platform (Illumina) in the rapid run mode (50 nt single-end reads). The list of all analysed smDNA libraries is shown in Supplementary Table S4.

### Genomic DNA library construction and sequencing

Genomic DNA was extracted from exponentially grown *E. coli* cells harvested at OD_600_=1.0 according to the published protocol ^35^. Preparation of genomic DNA libraries was carried out using the NEBNext Ultra II FS DNA Library Prep Kit (NEB) according to the manufacturer’s instructions. The approximate insert size was selected to be in the range of 150-250 bp. Barcodes were introduced to both ends during library amplification step with NEBNext multiplex oligos for Illumina (NEB). Genomic DNA libraries were sequenced using HiSeq2500 platform (Illumina) in the rapid run mode (50 nt single end reads). The list of all analysed genomic DNA libraries is shown in Supplementary Table S4.

### Analysis of high-throughput sequencing data

All libraries were quality checked before further processing using FastQC (v. 0.11.8). The 3’-adaptor sequence (5’-TGGAATTCTCGGGTGCCAAGGC-3’) was trimmed and reads with length less than 14 nt were removed using cutadapt (v. 2.7). Reads were aligned onto the reference genomes (Refseq accession numbers: NC_012971.2 in the case of BL21(DE3) and its derivatives, NC_000913.3 in the case of MG1655 and its derivatives, and GSE107973 in the case of strains containing the I-SceI site) and corresponding plasmids allowing zero mismatches via bowtie (v. 1.2.3). Genome coverage in the 1 Kb windows (RPKM) for uniquely mapped reads (total or strand-specific) were calculated using bedtools (v. 2.28.0). Multi-mapping reads that mapped to both plasmid and chromosome loci were filtered out, in order to avoid biases caused by different plasmid copy numbers. Genome coverage for chromosomal multi-mappers were calculated as described above with the exception that the value was divided by the number of read’s locations to which it was mapped. The chromosomal region between 2 and 3 Mb, which lacked prominent hotspots of smDNA loading, was used to analyse average smDNA distribution around Chi sites. In this case, the number of reads that mapped to the DNA strand with the Chi site in the 5’-GCTGGTGG-3’ orientation was calculated in 500 nt bins in the 20 Kb region centered around each Chi site and averaged.

The expected proportion of smDNA reads mapped to a plasmid was calculated according to the random sampling model as follows: proportion of reads = total number of reads×plasmid length/(genome length + plasmid length). The plasmid copy number was estimated based on information available from the literature (12 for pBAD, 15 for pChi and pNonChi, 20 for pET).

Nucleotide Logos (Fig. S1C and D) were generated using custom python scripts and WebLogo3 (v. 3.7). Only reads with the length of 18 nt were taken into analysis.

The significance of the IS1 and IS3 enrichment in the regions of preferential smDNA generation by CbAgo (Fig. S3) was analysed using a permutation test. For this purpose, 10,000 samples of 29 IS1 elements (11 in the case of IS3) randomly distributed across the *E. coli* genome were generated and intersected with the list of all genomic regions where the ratio of smDNA (CbAgo/dCbAgo) is greater than 1.

To analyse strand-specific distribution of smDNAs along the *E. coli* chromosome (Fig. 2F), the ratio of smDNAs that mapped to the plus- and minus-strands in 1 Kb windows was calculated and plotted against the genomic coordinate. A rolling mean (10 Kb window, 1 Kb step) was added.

Genomic DNA libraries were processed essentially as described above. Genome coverage in 1 Kb windows was calculated, smoothed with Loess regression and plotted against the chromosome coordinate.

All plots were generated in R (v. 3.6.1) using custom scripts.

### Determination of the efficiency of plasmid loss

For experiments on plasmid loss, *E. coli* MG1655Z1 strains containing chromosomal CbAgo/dCbAgo or lacking CbAgo (Table S2) were transformed with plasmids from various incompatibility groups (Table S1). Several colonies of transformed cell cultures from LB plates were inoculated into liquid LB medium with the addition of anhydrotetracycline (aTc) (0.1 μg/ml) and grown at 25°C until OD_600_~0.5. Then, glycerol was added to the cell culture to the final concentration of 20%, 200 μl aliquots were frozen in liquid nitrogen and stored at −70 °C.

Aliquots of frozen cells of *E. coli* MG1655Z1::CbAgo, MG1655Z1::dCbAgo or control MG1655Z1 were thawed in ice and inoculated into liquid LB with the addition of aTc and antibiotic corresponding to the plasmid used (Supplementary Table 2). The cell culture was grown at 25 °C and passaged in LB with the addition of aTc and without any antibiotics every 12 hours. The percentage of antibiotic resistant cells containing the plasmid was determined after 5 (for RSF1010 and pSRKTc), 8 (for pET28b), and 9 (for pBR325 and pACYC) passages by plating the cultures on agar media with and without antibiotic.

### Analysis of phage infection

#### Preparation of M13 phage lysate

Overnight culture of *E. coli* NEB Turbo strain (containing the F’ factor required for M13 infection) was diluted 10-fold with 5 ml of LB and a slice of agar with a plaque of phage M13 was added. The culture was incubated for 4-5 hours at 37 °C with aeration. The cells were precipitated by centrifugation (4000 g, 10 min) and the supernatant was filtered through a sterile syringe filter Millex-GP with a 0.22 µm hydrophilic Polyethersulfone membrane (Merсk, Millipore). The resulting lysate was stored at +4 °C for no more than one week.

#### Titration of phage M13

Night culture of *E. coli* NEB Turbo was grown in liquid LB, diluted 100-fold with fresh LB and grown at 37 °C with aeration until OD~0.5. A series of dilutions of bacteriophage lysate in liquid LB were prepared, and 10 μl of the lysate was mixed with 200 μl of cell cultures. The mixture was incubated at room temperature for 1-5 minutes to adsorb bacteriophage particles on the cells, mixed with 3 ml of molten (47 °C) 0.5% bacterial agar, poured into a Petri dish with solidified lower LB agar, and evenly distributed over the dish. The plates were incubated at 37 °C for 12-16 hours, and the number of negative colonies was counted.

#### Effects of CbAgo on phage production

For experiments on phage infection, cultures of *E. coli* NEB Turbo Z1::CbAgo, Z1::dCbAgo, or control strain without CbAgo (Supplementary Table S2) were grown and aliquoted as described above for experiments on plasmid loss. Aliquotes of these cultures were added to LB with aTc (0.1 μg/ml) and grown overnight at 25 °C. 200 µl of overnight cultures were inoculated into 20 ml of fresh LB+aTc, 10 µl of phage M13 lysate with the titer of 10^12^ was added, and grown at 25 °C with aeration. 5 ml aliquotes were taken after 2, 4, 6, and 8 hours and centrifuged (4000 g; 10 min). The supernatant was filtered through 0.22 μm filters. Each sample was titrated in the standard way using the *E. coli* NEB Turbo strain. For each strain, four biological replicates of the experiment were conducted.

### Analysis of the distribution of pAgos, RecBCD and AddAB in prokaryotic genomes

The proteins belonging to the RecBCD and AddAB systems were searched for in the NCBI protein database (downloaded in Dec 2019) using *hmmsearch* (v. 3.2.1) and HMM profiles from the TIGR database and Ref.^36^: RecB (HMM profile RecB, TIGR00609); RecC (recC, TIGR01450); RecD (recD1, TIGR01447); AddA (addA_alphas, TIGR02784; addA_Gpos, TIGR02785; AddA_cremie and AddA_epsilon, Ref. ^36^); AddB (addB_alphas, TIGR02786; addB_Gpos, TIGR02773; rexB_recomb; TIGR02774; AddB_cremie and AddB_epsilon, Ref. ^36^)

The protein was marked as the DSBR protein if it had a match with any of these HMM profiles (E-value ≤ 1e-3). If the protein had matches with several HMM profiles, then the match with the best score was selected. Identified DSBR proteins were attributed to NCBI genomic assemblies of prokaryotic strains (downloaded from NCBI FTP in Jan 2020), and then only 19,253 assemblies with “Complete Genome” or “Chromosome” statuses were further considered. The genome was considered as encoding RecBCD or AddAB systems if all proteins from the corresponding system were identified as encoded in this genome.

The search of pAgo proteins in the recent NCBI protein database (downloaded in Dec 2019), their alignment, phylogenetic analysis and classification were performed as in Ref.^2^. Some strains have several NCBI accessible genomic assemblies, and in the order to remove such redundancy only the largest genomic assembly with the largest number of annotated proteins was considered for each strain.

## Data and Code Availability

All data generated during this study are included in the published article and the Extended Data and are available from public databases. The codes used for data analysis are available from the corresponding authors upon reasonable request.

## Acknowledgements

We thank Dr. Martin A. White for strains with engineered I-SceI sites, Dr. Gerry Smith for RecBCD antibodies. The project was in part supported by the Ministry of Science and Higher Education of the Russian Federation (14.W03.31.0007), Russian Science Foundation (19-14-00359), Russian Foundation for Basic Research (18-29-07086). DL is supported by grant MR/M019160/1 from the MRC (UK).

---

**Extended Data Fig. 1.**
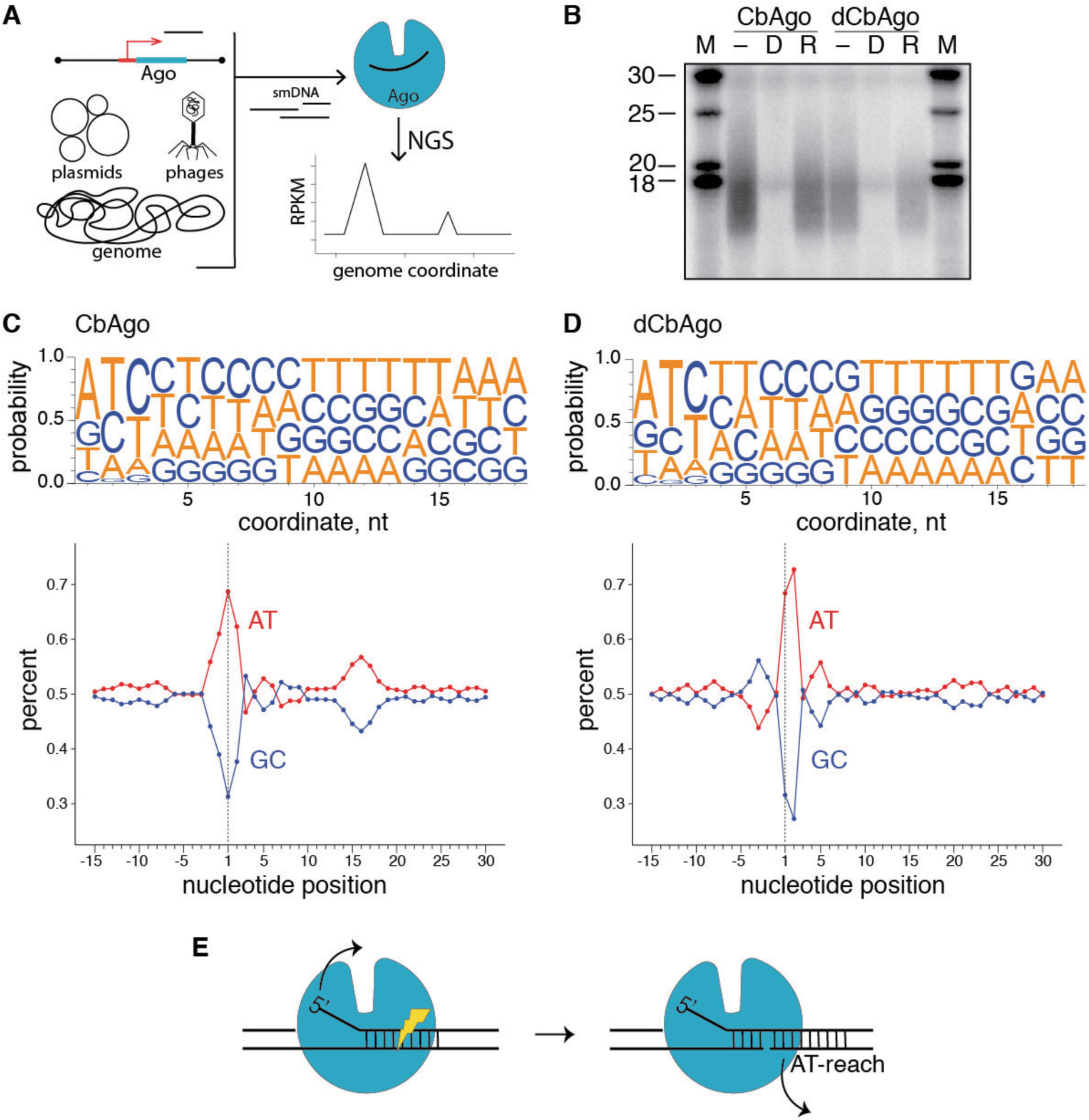
Small DNAs associated with CbAgo. (a) Scheme of experiments. The CbAgo gene with plasmid or chromosomal locations is expressed from an inducible promoter, followed by isolation of CbAgo-associated smDNAs, their sequencing and mapping to genomic coordinates of the chromosome, plasmids and phages. (b) Analysis of small nucleic acids isolated from wild-type CbAgo and dCbAgo. The samples were treated with alkaline phosphatase, ^32^P-labelled with polynucleotide kinase and treated with DNaseI (D), RNase (R) or left without further treatment (-). The DNA marker (M) lengths are indicated. (c) and (d) Analysis of nucleotide biases for smDNA guides for wild-type CbAgo (c) and dCbAgo (d). (*Top*) Nucleotide frequencies at different guide positions. (*Bottom*) AT/GC-content along the guide length and in surrounding genomic sequences. Guide positions starting from the 5’-end are indicated below the plots. The AT-bias around the first position is seen for both active CbAgo and dCbAgo. The AT-bias in the downstream region (positions 14-18) is seen for active CbAgo but not for dCbAgo. (e) Model of processing of smDNAs by CbAgo from double-stranded DNA precursors. Binding of the guide 5’-end in the MID-pocket of CbAgo may be facilitated by melting of the DNA duplex in the upstream guide region (left scheme). Guide DNA loading is completed after CbAgo-dependent cleavage of the complementary DNA strand and its dissociation, depending on the AT-richness of the downstream guide-target duplex (right scheme).

**Extended Data Fig. 2.**
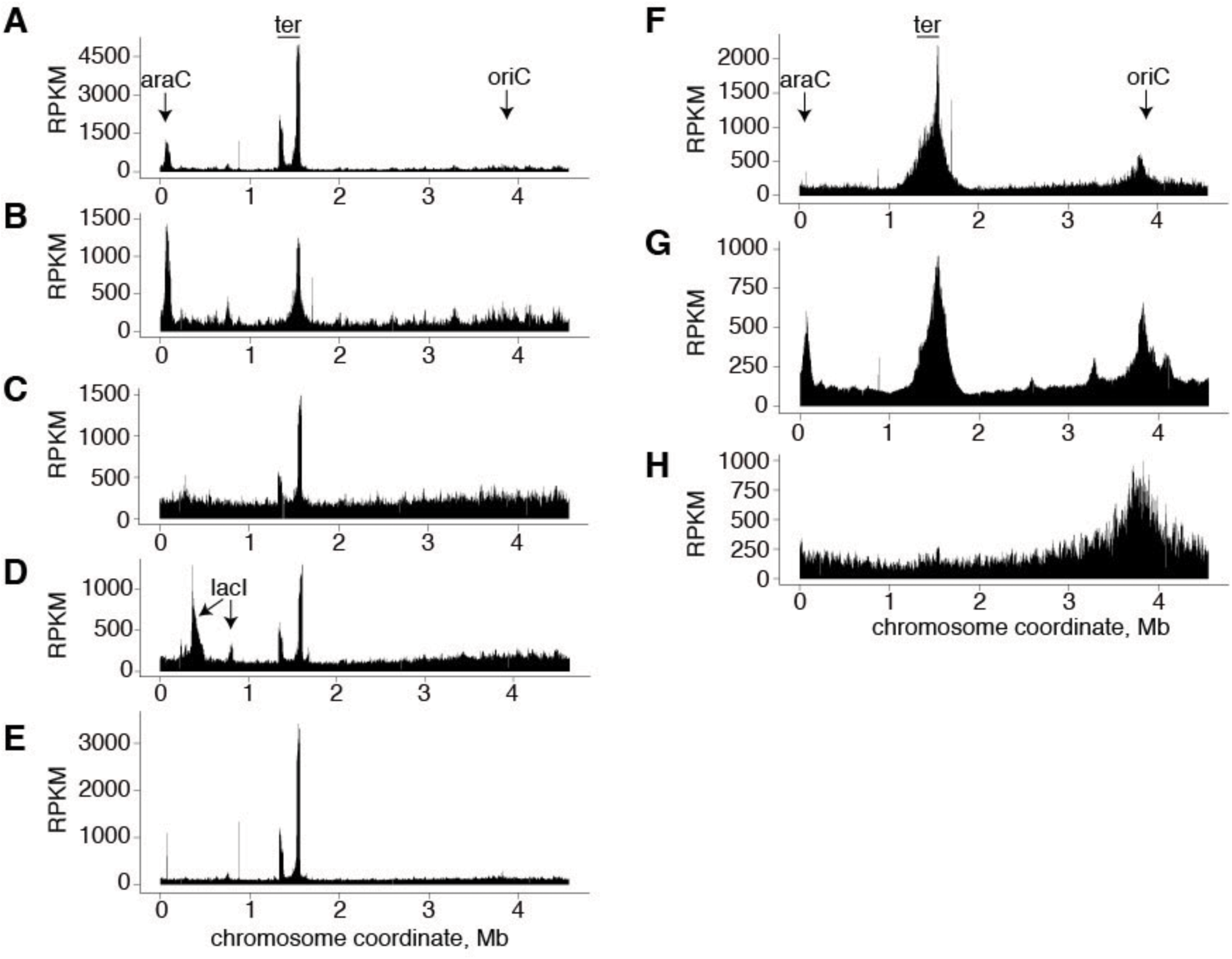
Whole-genome mapping of smDNAs in strains with various genetic backgrounds. For each strain, the distribution of smDNAs along the chromosome is shown in RPKM (the number of smDNAs reads per kilobase per million reads in the smDNA library). Positions of the *araC*, *lacI* and *Ter* sites are shown above the plots. The identities of the strains and plasmids, with plasmid or chromosomal localizations of the CbAgo gene, are indicated (see Supplementary Tables 1 and 2). (a) Wild-type CbAgo with plasmid localization (pBAD containing the *araC* gene) in BL21(DE3). (b) The same in BL21(DE3) with knockout of Tus. (c) Wild-type CbAgo with genomic localization in MG1655Z1. (d) The same with pET28b containing *lacI*. (e) Catalytically dead dCbAgo with the plasmid localization (pBAD) in BL21(DE3). (f) Wild-type CbAgo with plasmid localization in BL21(DE3) with knockout of RecB/RecD. (g) Wild-type CbAgo with plasmid localization in BL21(DE3) with knockout of RecC. (h) Wild-type CbAgo with plasmid localization in BL21(DE3) with knockout of RecA.

**Extended Data Fig. 3.**
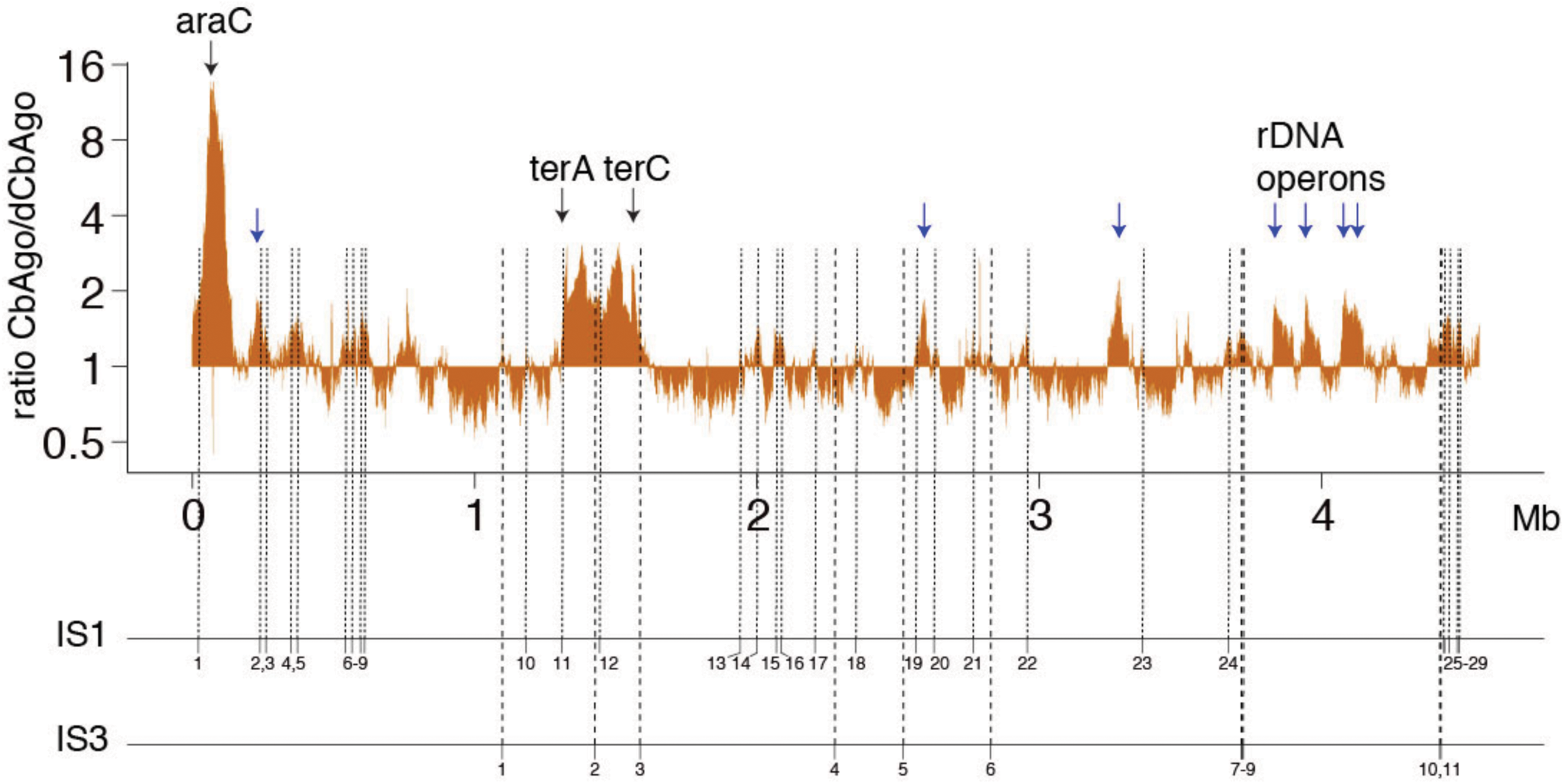
Targeting of specific genomic regions depends on the catalytic activity of CbAgo. The ratio of smDNAs for wild-type CbAgo and dCbAgo (obtained for the BL21(DE3) pBAD_CbAgo strains), shown in the logarithmic scale. Normalized densities of smDNA reads (RPKM in 1 kb bins) were calculated for each CbAgo variant and divided by each other. The regions with the ratio of >1 correspond to the sites of active smDNA processing by CbAgo. CbAgo targets the *araC* locus, *Ter* region and multicopy sequences: rDNA operons (indicated with arrows above the plot) and IS elements. Positions of IS1 (29 copies) and IS3 (12 copies) in the BL21(DE3) genome are shown with dotted lines below the plot.

**Extended Data Fig. 4.**
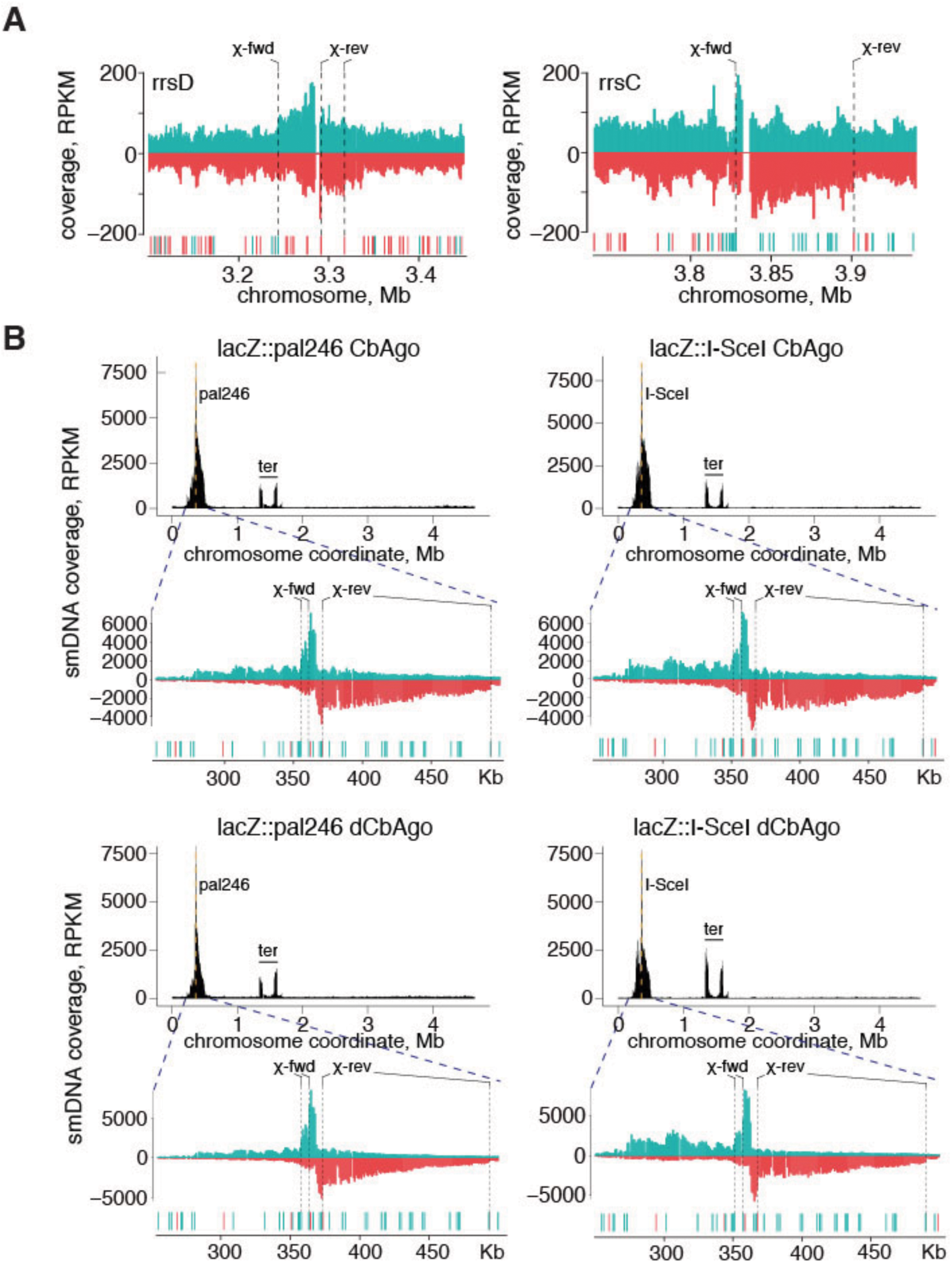
RecBCD-dependent asymmetry in smDNA distribution at multicopy regions and the sites of engineered DSBs. (a) Examples of smDNA distributions around rRNA operons, *rrsD* and *rrsC*. The reads from the plus and minus genomic strands are shown in green and red, respectively. Positions of Chi sites in surrounding genomic regions are indicated (forward for the plus strand and reverse for the minus strand); the closest Chi sites in the proper orientation are shown with dotted lines. (b) SmDNA peaks around engineered DSBs (palindrome- or I-SceI-dependent) for wild-type CbAgo (*top*) or dCbAgo (*bottom*). Most smDNAs are produced from the 3’-strand at each end of the DSB, and the boundaries of the smDNA peaks depend on Chi sites.

**Extended Data Fig. 5.**
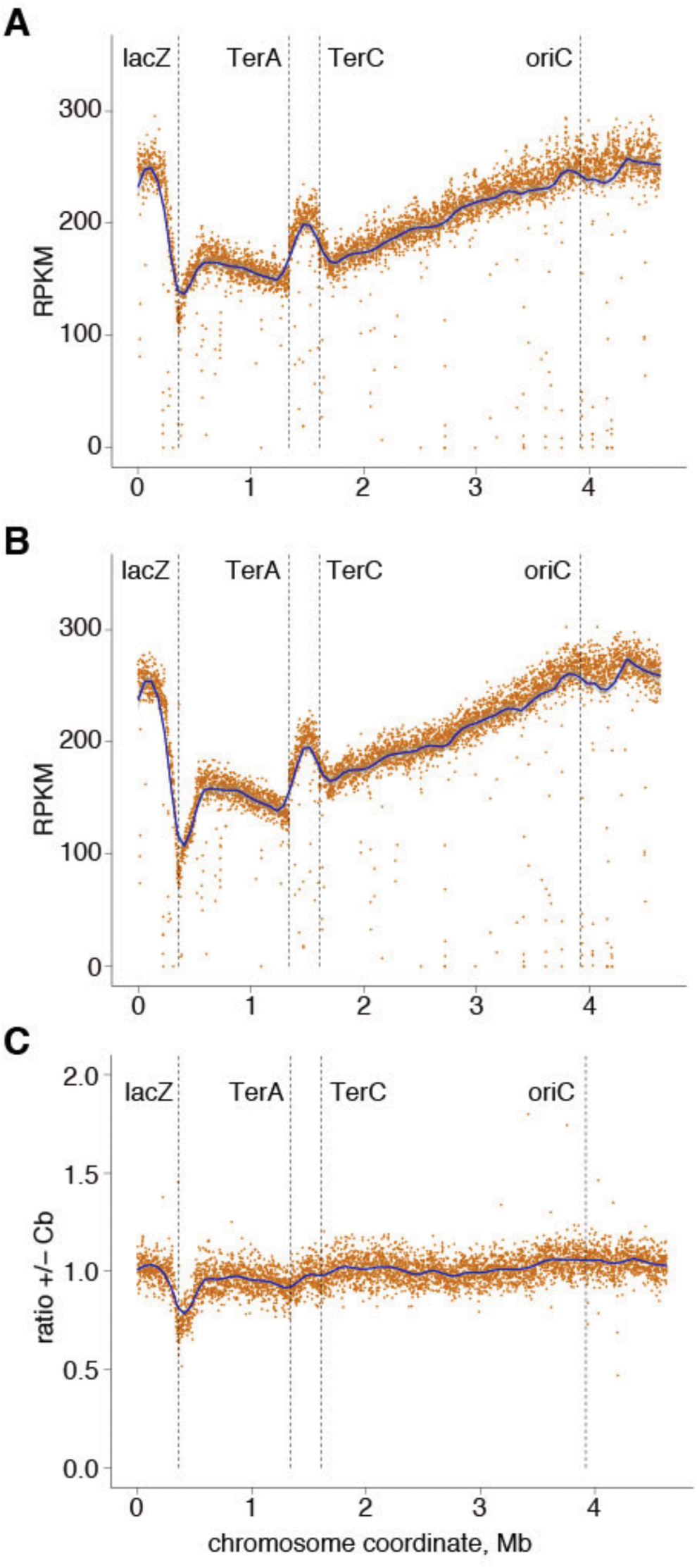
Genomic DNA coverage at DSBs formed by the I-SceI meganuclease. (a) *E. coli* strain with induced I-SceI but without expression of CbAgo. (b) *E. coli* strain with expression of both I-SceI and CbAgo. (c) The ratio of genomic DNA profiles for the strains with and without expression of CbAgo. Genomic DNA coverage is shown in RPKM.

**Extended Data Fig. 6.**
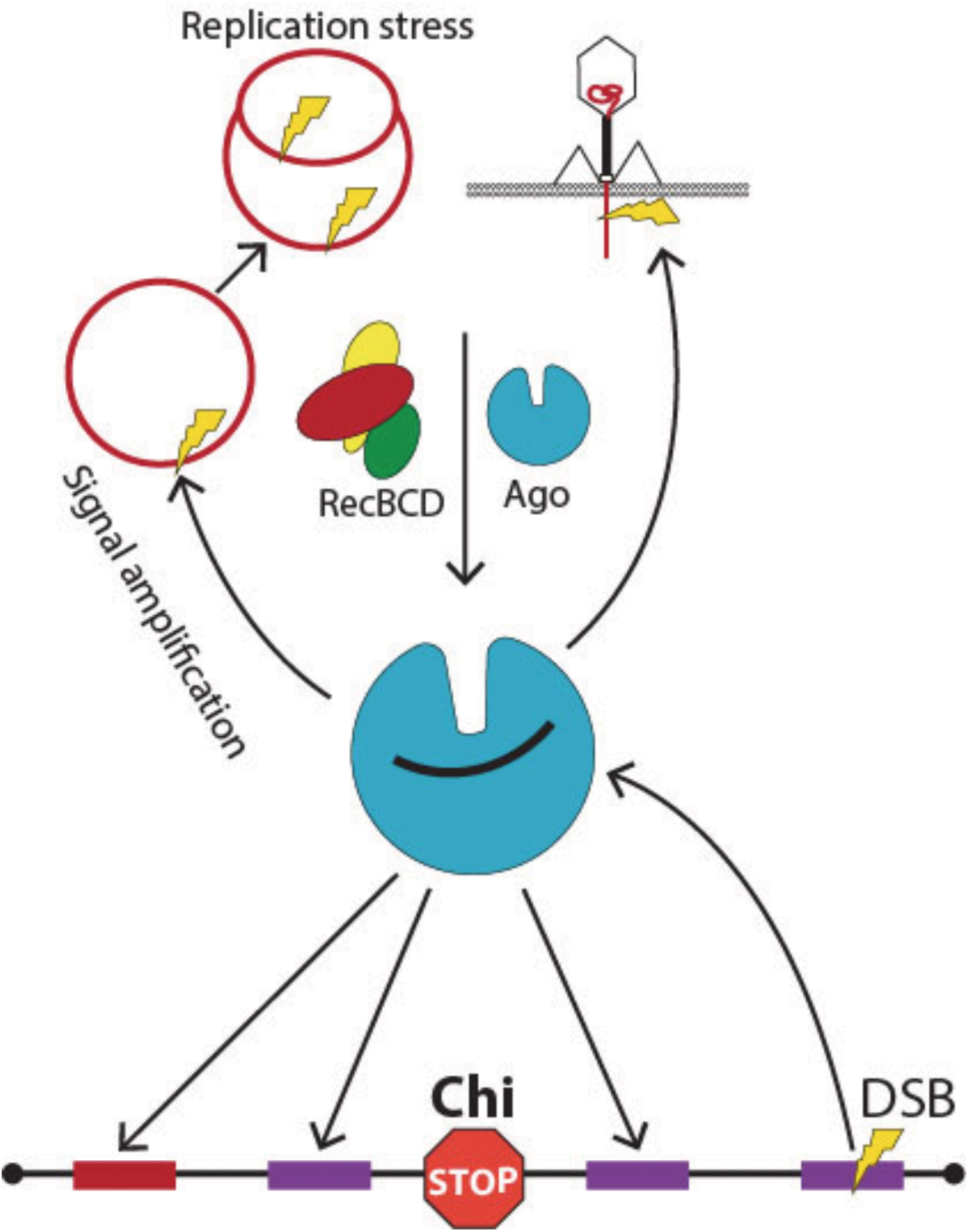
The proposed mechanism of DNA interference by CbAgo. Invader genetic elements, such as plasmids and phages, are actively processed by CbAgo and RecBCD due to their multicopy nature, intense replication, and the absence of Chi sites. This results in preferential loading of small guide DNAs corresponding to these elements into CbAgo and their directed cleavage leading to signal amplification. CbAgo can also attack chromosomal loci with regions of homology to plasmids (red) and multicopy genomic elements (violet), but extensive chromosome degradation is stopped due to the presence of Chi sites in the host DNA.

**Extended Data Fig. 7.**
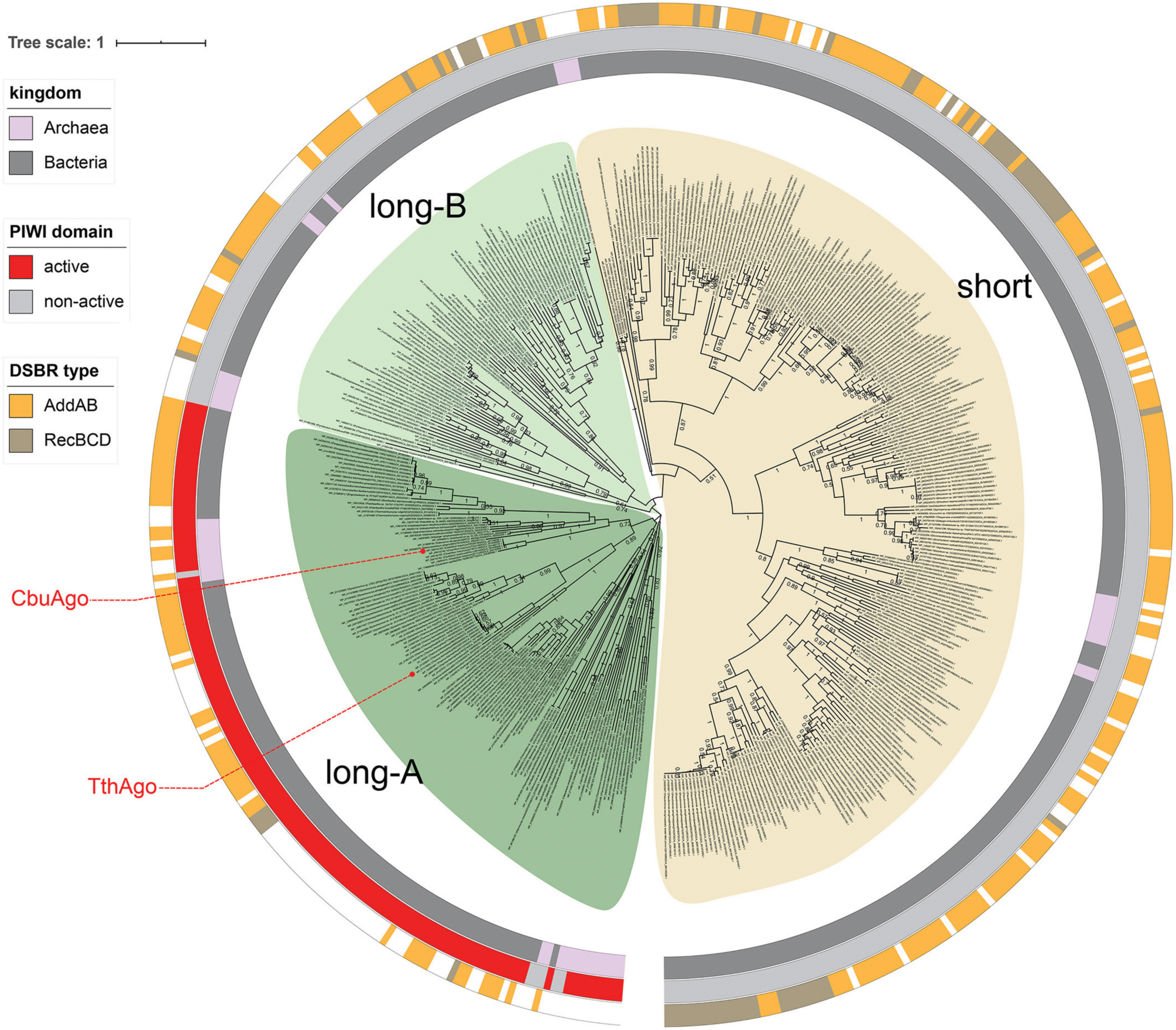
Co-occurrence of pAgo proteins and DSB repair systems in prokaryotic genomes. The circular phylogenetic tree of pAgos from bacterial strains with fully assembled genomes was constructed based on the multiple alignment of the MID-PIWI domains. The pAgo proteins were annotated as follows, from the inner to the outer circles: the superkingdom to which the corresponding pAgo belongs; the type of the PIWI domain, depending on the presence of the catalytic tetrad DEDX; the type of the DSB repair system encoded in the corresponding genome. CbAgo and TtAgo are highlighted in red. The scale bar represents the evolutionary rate calculated under the JTT+CAT evolutionary model. RecBCD and AddAB are encoded in 7,419 (38.5%) and 8,141 (42.3%) prokaryotic genomic assemblies, correspondingly. Both RecBCD and AddAB are found in 56 (0.3%) genomes. In total, we found 1,711 pAgos encoded in 2,883 genomes of 2,802 bacterial strains, including 833 pAgos that were not found previously^2^. For the construction of the PIWI-MID-based phylogenetic tree we selected a subset of 399 pAgos of 456 strains with fully sequenced genomes (“Complete Genome” or “Chromosome” statuses).

## References

1. Makarova, K. S., Wolf, Y. I., van der Oost, J. & Koonin, E. V. Prokaryotic homologs of Argonaute proteins are predicted to function as key components of a novel system of defense against mobile genetic elements. Biology direct 4, 29, doi:10.1186/1745-6150-4-29 (2009).

2. Ryazansky, S., Kulbachinskiy, A. & Aravin, A. A. The Expanded Universe of Prokaryotic Argonaute Proteins. mBio 9, e01935–01918, doi:10.1128/mBio.01935-18 (2018).

3. Swarts, D. C. et al. The evolutionary journey of Argonaute proteins. Nature structural & molecular biology 21, 743–753, doi:10.1038/nsmb.2879 (2014).

4. Hegge, J. W. et al. DNA-guided DNA cleavage at moderate temperatures by Clostridium butyricum Argonaute. Nucleic acids research 47, 5809–5821, doi:10.1093/nar/gkz306 (2019).

5. Kuzmenko, A., Yudin, D., Ryazansky, S., Kulbachinskiy, A. & Aravin, A. A. Programmable DNA cleavage by Ago nucleases from mesophilic bacteria Clostridium butyricum and Limnothrix rosea. Nucleic acids research 47, 5822–5836, doi:10.1093/nar/gkz379 (2019).

6. Swarts, D. C. et al. Argonaute of the archaeon Pyrococcus furiosus is a DNA-guided nuclease that targets cognate DNA. Nucleic acids research 43, 5120–5129, doi:10.1093/nar/gkv415 (2015).

7. Swarts, D. C. et al. DNA-guided DNA interference by a prokaryotic Argonaute. Nature 507, 258–261, doi:10.1038/nature12971 (2014).

8. Zander, A. et al. Guide-independent DNA cleavage by archaeal Argonaute from Methanocaldococcus jannaschii. Nature microbiology 2, 17034, doi:10.1038/nmicrobiol.2017.34 (2017).

9. Willkomm, S., Makarova, K. & Grohmann, D. DNA-silencing by prokaryotic Argonaute proteins adds a new layer of defence against invading nucleic acids. FEMS microbiology reviews 42, 376–387, doi:10.1093/femsre/fuy010 (2018).

10. Lisitskaya, L., Aravin, A. A. & Kulbachinskiy, A. DNA interference and beyond: structure and functions of prokaryotic Argonaute proteins. Nature communications 9, 5165, doi:10.1038/s41467-018-07449-7 (2018).

11. Gebert, D. & Rosenkranz, D. RNA-based regulation of transposon expression. Wiley interdisciplinary reviews. RNA 6, 687–708, doi:10.1002/wrna.1310 (2015).

12. Meister, G. Argonaute proteins: functional insights and emerging roles. Nat Rev Genet 14, 447–459, doi:10.1038/nrg3462 (2013).

13. Kaya, E. et al. A bacterial Argonaute with noncanonical guide RNA specificity. Proceedings of the National Academy of Sciences of the United States of America 113, 4057–4062, doi:10.1073/pnas.1524385113 (2016).

14. Sheng, G. et al. Structure-based cleavage mechanism of Thermus thermophilus Argonaute DNA guide strand-mediated DNA target cleavage. Proceedings of the National Academy of Sciences of the United States of America 111, 652–657, doi:10.1073/pnas.1321032111 (2014).

15. Willkomm, S. et al. Structural and mechanistic insights into an archaeal DNA-guided Argonaute protein. Nature microbiology 2, 17035, doi:10.1038/nmicrobiol.2017.35 (2017).

16. Olina, A. et al. Genome-wide DNA sampling by Ago nuclease from the cyanobacterium Synechococcus elongatus. RNA biology, 1–12, doi:10.1080/15476286.2020.1724716 (2020).

17. Swarts, D. C. et al. Autonomous Generation and Loading of DNA Guides by Bacterial Argonaute. Molecular cell 65, 985–998, doi:10.1016/j.molcel.2017.01.033 (2017).

18. Olovnikov, I., Chan, K., Sachidanandam, R., Newman, D. K. & Aravin, A. A. Bacterial argonaute samples the transcriptome to identify foreign DNA. Molecular cell 51, 594–605, doi:10.1016/j.molcel.2013.08.014 (2013).

19. Duggin, I. G. & Bell, S. D. Termination structures in the Escherichia coli chromosome replication fork trap. Journal of molecular biology 387, 532–539, doi:10.1016/j.jmb.2009.02.027 (2009).

20. Dillingham, M. S. & Kowalczykowski, S. C. RecBCD enzyme and the repair of double-stranded DNA breaks. Microbiology and molecular biology reviews: MMBR 72, 642–671, Table of Contents, doi:10.1128/MMBR.00020-08 (2008).

21. Smith, G. R. How RecBCD enzyme and Chi promote DNA break repair and recombination: a molecular biologist’s view. Microbiology and molecular biology reviews: MMBR 76, 217–228, doi:10.1128/MMBR.05026-11 (2012).

22. Wigley, D. B. Bacterial DNA repair: recent insights into the mechanism of RecBCD, AddAB and AdnAB. Nature reviews. Microbiology 11, 9–13, doi:10.1038/nrmicro2917 (2013).

23. Sinha, A. K. et al. Division-induced DNA double strand breaks in the chromosome terminus region of Escherichia coli lacking RecBCD DNA repair enzyme. PLoS genetics 13, e1006895, doi:10.1371/journal.pgen.1006895 (2017).

24. Sinha, A. K. et al. Broken replication forks trigger heritable DNA breaks in the terminus of a circular chromosome. PLoS genetics 14, e1007256, doi:10.1371/journal.pgen.1007256 (2018).

25. White, M. A., Azeroglu, B., Lopez-Vernaza, M. A., Hasan, A. M. M. & Leach, D. R. F. RecBCD coordinates repair of two ends at a DNA double-strand break, preventing aberrant chromosome amplification. Nucleic acids research 46, 6670–6682, doi:10.1093/nar/gky463 (2018).

26. Eykelenboom, J. K., Blackwood, J. K., Okely, E. & Leach, D. R. SbcCD causes a double-strand break at a DNA palindrome in the Escherichia coli chromosome. Molecular cell 29, 644–651, doi:10.1016/j.molcel.2007.12.020 (2008).

27. White, M. A., Darmon, E., Lopez-Vernaza, M. A. & Leach, D. R. F. DNA double strand break repair in Escherichia coli perturbs cell division and chromosome dynamics. PLoS genetics 16, e1008473, doi:10.1371/journal.pgen.1008473 (2020).

28. El Karoui, M., Biaudet, V., Schbath, S. & Gruss, A. Characteristics of Chi distribution on different bacterial genomes. Research in microbiology 150, 579–587, doi:10.1016/s0923-2508(99)00132-1 (1999).

29. Jolly, S. M. et al. A DNA-guided Argonaute Protein Functions in DNA Replication in Thermus thermophilus. bioRxiv, 869172, doi:https://doi.org/10.1101/869172 (2019).

30. Koonin, E. V. Evolution of RNA- and DNA-guided antivirus defense systems in prokaryotes and eukaryotes: common ancestry vs convergence. Biology direct 12, 5, doi:10.1186/s13062-017-0177-2 (2017).

31. Modell, J. W., Jiang, W. & Marraffini, L. A. CRISPR-Cas systems exploit viral DNA injection to establish and maintain adaptive immunity. Nature 544, 101–104, doi:10.1038/nature21719 (2017).

32. Levy, A. et al. CRISPR adaptation biases explain preference for acquisition of foreign DNA. Nature 520, 505–510, doi:10.1038/nature14302 (2015).

33. Datsenko, K. A. & Wanner, B. L. One-step inactivation of chromosomal genes in Escherichia coli K-12 using PCR products. Proceedings of the National Academy of Sciences of the United States of America 97, 6640–6645, doi:10.1073/pnas.120163297 (2000).

34. Bohn, C., Collier, J. & Bouloc, P. Dispensable PDZ domain of Escherichia coli YaeL essential protease. Molecular microbiology 52, 427–435, doi:10.1111/j.1365-2958.2004.03985.x (2004).

35. He, F. E. coli Genomic DNA Extraction Bio-101, e97, doi:10.21769/BioProtoc.97 (2011).

36. Bernheim, A., Bikard, D., Touchon, M. & Rocha, E. P. C. A matter of background: DNA repair pathways as a possible cause for the sparse distribution of CRISPR-Cas systems in bacteria. Philosophical transactions of the Royal Society of London. Series B, Biological sciences 374, 20180088, doi:10.1098/rstb.2018.0088 (2019).

